# Early warning signs in social-ecological networks

**DOI:** 10.1101/003269

**Authors:** Samir Suweis, Paolo D'Odorico

## Abstract

A number of social-ecological systems exhibit complex behaviour associated with nonlinearities, bifurcations, and interaction with stochastic drivers. These systems are often prone to abrupt and unexpected instabilities and state shifts that emerge as a discontinuous response to gradual changes in environmental drivers. Predicting such behaviours is crucial to the prevention of or preparation for unwanted regime shifts. Recent research in ecology has investigated early warning signs that anticipate the divergence of univariate ecosystem dynamics from a stable attractor. To date, leading indicators of instability in systems with multiple interacting components have remained poorly investigated. This is a major limitation in the understanding of the dynamics of complex social-ecological networks. Here, we develop a theoretical framework to demonstrate that rising variance – measured, for example, by the maximum element of the covariance matrix of the network – is an effective leading indicator of network instability. We show that its reliability and robustness depend more on the sign of the interactions within the network than the network structure or noise intensity. Mutualistic, scale free and small world networks are less stable than their antagonistic or random counterparts but their instability is more reliably predicted by this leading indicator. These results provide new advances in multidimensional early warning analysis and offer a framework to evaluate the resilience of social-ecological networks.

## Introduction

Social-ecological systems are often difficult to investigate and manage because of their inherent complexity (1). Small variations in external drivers can lead to abrupt changes associated with instabilities and bifurcations in the underlying dynamics (2–4). These transitions can occur in a variety of ecological and social systems, and are often unexpected and difficult to revert (4). Anticipating critical transitions and divergence from the present state of the system is particularly crucial to the prevention or mitigation of the effects of unwanted and irreversible changes (5–10). Recent research in ecology has focused on leading indicators of regime shift in ecosystems characterized by one state variable (5,7,11,12). These indicators are typically associated with the critical slowing down phenomenon: as the system approaches a critical transition, its response to small perturbations of the stable state becomes slower (11). It has been shown that in univariate systems (i.e., with only one state variable) critical slowing down entails an increase in the temporal variance and autocorrelation of the state variable (5). The case of systems with several mutually interacting components, however, has remained poorly investigated (13–15), while the connection between network stability and research on indicators for loss of resilience has been elusive (16).

Here we develop a theoretical framework to analyze early warning signs of instability and regime shift in complex networks. We provide analytical expressions for a set of precursors of instability in complex systems with additive noise for a variety of network structures.

We consider a social-ecological system with *N* components (nodes) coupled through a set of links. The state of the system is expressed by the vector ***x*** of length *N*, whose terms *x_i_* represent the state of node *i*. The local stability of a state ***x\**** is evaluated through a linearization, 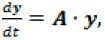 where **y** = **x**-**x\*** is the displacement of **x** from **x\***; **A** is the *N* × *N* matrix expressing the interactions among nodes in the (linearized) dynamics (see Methods). In population ecology this framework is typically used to express the dynamics of a community of *N* populations interacting according to the relationships determined by the matrix **A**, often known as “*community matrix*”(2,17–19); likewise, in social systems **A** describes the network of interactions (e.g., trade, migration, flow of information among people, groups of individuals, or countries (20–23)). The off-diagonal terms of **A** determine the pairs of interacting nodes as well as the strength of their interaction. The dynamics are stable if the maximum real part of the eigenvalues of **A**, *Max*[Re(λ)], is negative.

Classic ecological theories (2,3) have considered the case of networks with randomly connected nodes (with a certain probability, *C*). The strength (*p*) of the interactions between them is represented by a zero-mean random variable of variance σ^2^. May (2,3) showed that random networks become unstable as connectivity (i.e., *C*), size (i.e., *N*) or the strength variance increase. The stability of networks with prescribed architectures (e.g., predator-prey, competitive or mutualistic interactions) also depends on connectivity, strength variance, system size, as well as on the network structure (17,19).

More generally, the off-diagonal terms of **A** may result from a set of “rules” expressed as a function of a few parameters of which connectivity and strength variance are just an example. Changes in the structure and intensity of the interactions correspond to variations in these parameters, which, in turn, can lead to instability by modifying the community matrix and its eigenvalues. How can we evaluate whether ongoing changes in the interactions within a socialecological network are reducing its resilience? Is there a way to use measurable quantities to determine whether the system is about to become unstable?

In one-dimensional systems leading indicators are typically associated with behaviors resulting from the eigenvalue tending to zero at the onset of instability. This effect entails a slower return to equilibrium after a “small” perturbation (11,24). Known as “critical slowing down”, this phenomenon exists also in systems with multiple interacting components, though it is hard to recognize and therefore it does not constitute an effective leading indicator of instability. In fact, in “real world” applications the equations driving the dynamics are not known and, therefore, the network nodes in which slowing down is expected to occur are not known a priori. Critical slowing down, however, has been related to an increase in variance and autocorrelation in the state variable of one dimensional systems (5,7,25). Here we provide a theoretical framework to investigate early-warnings in the variance, autocorrelation, and power spectrum of multidimensional systems with interactions described by a given network structure.

### Methods

We consider a network with *N* interacting nodes. The state of the system, **x***=*{*x*_1_, *x*_2_, … *x*_N_}, is governed by dynamics: d***x*** = *f*(*x*,*p*,*C*)d*t* + *v* **I** d*W*, where ***f***={*f*_1_, *f*_2_, …, *f*_N_} is a *N*-dimensional vector function expressing the deterministic component of the dynamics of ***x***, as a function of a set of parameters, *p* and *C*; **I** is the identity matrix, and *ν* d*W* is an additive stochastic driver represented by a white Gaussian noise of mean zero and intensity *ν*d*t*. If we consider a small perturbation **y** forcing the system away from its equilibrium point **x\*** (i.e., **y** = **x**−**x\***), inserting **x** = **x\***+**y** in the above equation and linearizing ***f***(**x\***+**y**, *p*, *C*) around **x\*** we obtain

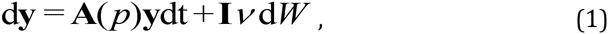

where 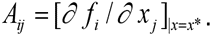 Eq. (1) is a multivariate Ornstein–Uhlenbeck process (26).

The stable states, **x\***, of Eq. (1) are the same as those of their deterministic counterparts, 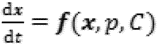 (27). These states are stable if the maximum real part of the eigenvalues of **A** is negative. To identify early warning signs of network instability, we relate the steady state covariance matrix 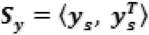 to the eigenvalues, λ, of **A**, where **y**_s_ is calculated from the steady state solution of Eq. (1). We first look for leading indicators of instability in the behavior of the covariance matrix, **S_y_**, of **y** as the system approaches instability. The (*i*,*j*) element of ***S*_y_** is: 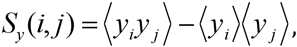 where 〈 〉 represents the average. The covariance matrix of the stationary dynamics of the system can be obtained (26) as the solution of Eq. (2):

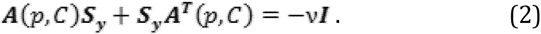

***S*_y_** is a function of the linearization matrix, ***A***(*p,C*), which, in turn, depends on the control parameters (*p or C*). At the onset of instability (i.e., as Max[Re(λ)] →0) the maximum element of the covariance matrix, **S_y_**, of **y** increases. More details on the time-lag correlation and power spectrum can be found in the Supplementary Materials. While the linearization matrix, ***A***, here accounts the interconnections existing among nodes within the network (i.e. the pairs of nodes that are connected by a link (3, 17)), the covariance matrix, **S_y_**, expresses the variance of the fluctuations of the state variable at each node (diagonal terms) and the interrelationship (positive or negative) of the fluctuations between pairs of nodes (off-diagonal terms). To better understand the structure of **S_y_**, we look at the case of a network with only two nodes. In this case the above equation for the covariance matrix can be solved analytically, and the covariance matrix reads (26)

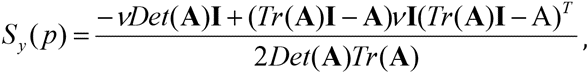

where *Det* is the determinant and *Tr* the trace of the matrix, which can be expressed as a function of the eigenvalues, *λ*_1,2_.

Thus, the covariance matrix diverges for λ_1_, λ_2_→0. The time correlation matrix, **ρ_y_**(Δ), can also be computed analytically and also diverges for λ_1_, λ_2_→0, independently of Δ. The general analytical expressions of **ρ_y_**(Δ) and of the power spectrum of **y** are also reported in the Supplementary Materials.

We generate networks of size *N*, with a variety of architectures for **A** (see Supplementary Materials), and reach instability either by keeping constant the connectivity, *C*, while changing the strength of the interactions, *p*, or by varying *C* for a fixed *p* (2,17,19). We then use the analytical relationship between the steady state covariance matrix, **S_y_**, of **y** and the eigenvalues of the matrix **A** (Eq. (2)). Similarly, we express the time-lag correlation, **ρ_y_**, and the power spectrum, **P_y_**, of **y** as a function of **A** and its eigenvalues.

## Results and discussion

We find that the elements of both **S_y_**and **ρ_y_** increase as the system approaches instability (i.e., *Max*[Re(λ)]→0). Therefore, we investigate potential indicators for early warning in the behavior of suitable components of **S_y_**, **ρ_y_** and **P_y_** for *Max*[Re(λ)]→0. To that end we first consider the components of **S_y_** corresponding to the most connected, the most central (28) and the least connected nodes of the network. We also consider indicators based on the properties of the entire network, such as the maximum and the difference between the maximum and minimum of the matrix **S_y_**.

Most of the indicators based on the covariance matrix, **S**_y_, have a non-trivial dependence on *Max*[Re(λ)] (see Figures 1, S1–S6). The maximum element of **S_y_** (*Max*[**S_y_**]) and *Max*[**S_y_**]-*Min*[**S_y_**] provide the most effective indicator of early warning in most networks (Figures 1, S7 and S8). In mutualistic (++) networks *Max*[**S_y_**] corresponds to the most connected node (the “hub”), regardless of their topological structure (Supplementary Materials, Figures S9–S10). All these indicators based on **S_y_** improve their performances when the size, *N*, of the network increases (compare main panels to insets in Figure 1; see also Supplementary Materials, Figure S11). Thus our ability to detect early warning signs and predict tipping points is enhanced in more diverse systems (16).

**Figure 1.**
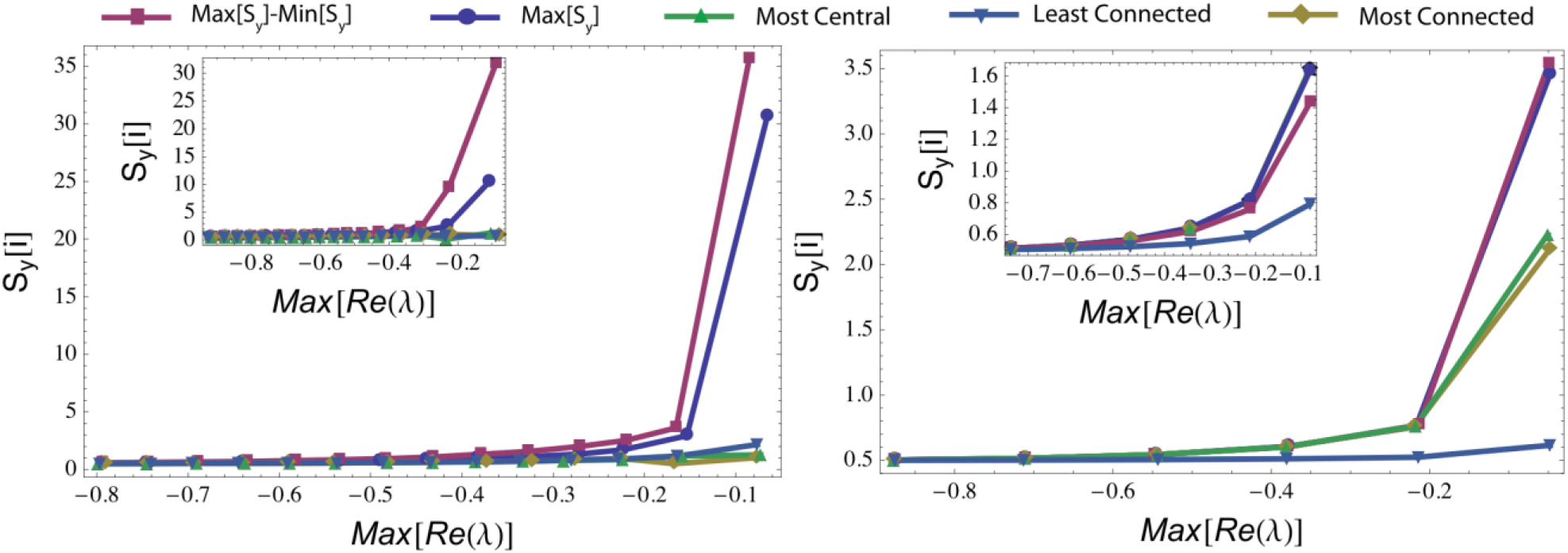
Leading indicators of instability based on different elements of the covariance matrix (**S_y_**), including the maximum (in absolute value) element, *Max*[**S_y_**], the difference between *Max*[**S_y_**] and *Min*[**S_y_**], the element of **S_y_** corresponding to the most connected, least connected, or highest eigenvector centrality (24) network node. Random (l*eft*) and scale free (right) (30) network generated with *N* = 50 and *C* = 0.1 (main panels) and *N* = 0.1 and *C* = 0.5 (insets). Instability (i.e., decrease in *Max*[*Re*(λ)]) is attained by increasing the interaction strength *p* (mean field case). The figures represent average behavior over 100 realizations.

We also look at the relationship between the maximum element of the time-lag correlation matrix, **ρ_y_**(Δ) (where Δ is the time lag), and *Max*[Re(λ)] for different values of Δ, *p* and *C* (Figures S12–S14). Although significant, these indicators are less efficient with respect to the case with zero time-lag (i.e., indicators based on **S_y_**). Finally, the power spectrum does not appear to be an effective indicator, as we identified only weak changes in **P_y_** for increasing values of *p* and *Max*[Re(λ)] (see Supplementary Materials, Figure S15). Therefore, here we focus on early warning signs provided by the way *Max*[**S_y_**] varies as a function of changes in *Max*[Re(λ)]. A warning sign is effective if (a) it appears in time to prevent (or prepare for) the occurrence of instability (29–30); (b) it relies on a well-defined and easy to recognize indicator (e.g., a detectable or significant increase in variance (29–30)); and (c) it does not give false positives (or false negatives) (31). We use these criteria to evaluate the effectiveness of *Max*[**S_y_**] as a leading indicator of instability with different network structures and levels of noise (32).

To investigate the effect of noise, we first consider the “mean-field” case of networks in which the absolute value of the interaction strength between connected nodes is a constant, *p*; we gradually increase *p* or *C* until *Max*[Re(λ)] becomes positive (17–19). We observe (Figure 2) a consistent increase in *Max*[***S*_y_**] for all network structures, regardless of whether instability is attained by increasing interaction strength or connectivity (Figures S17–S6). The network structure, however, affects the timeliness of *Max*[**S_y_**] as a leading indicator. In fact, *Max*[***S*_y_**] exhibits a more defined increase and a better anticipation of the onset of instability in the case of random networks than with all the other structures. In the case of these “mean field” networks we did not consider the antagonistic structure because antagonistic networks with constant interaction strength (in absolute value) are always stable regardless of the parameters *p* and *C* (see Supplementary Materials, Figs. S1, S2).

**Figure 2.**
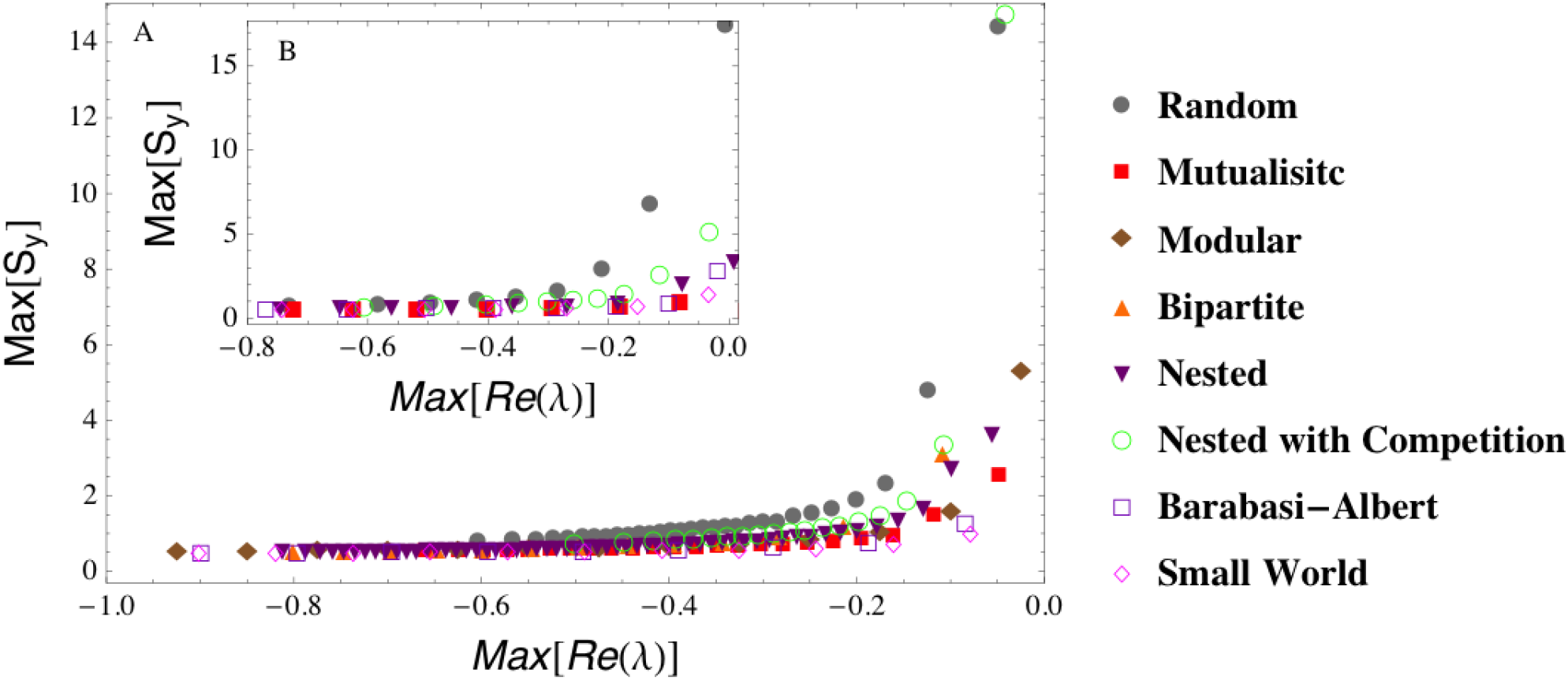
*Max*[**S_y_**]as a leading indicator of instability in a “mean field” network with constant interaction intensity (in absolute value), *p*. Instability is attained by increasing *p* (main panel A, with *N* = 20, *C* = 0.2) or *C* (inset B, with *N* = 20, and *C* increasing from 0.1 to 1) with different network structures. The figures represent average behavior over 1000 realizations.

Likewise, in the case of random interaction strengths *Max*[***S*_y_**] exhibits a well-defined increase and a better anticipation of the instability in random networks than with the more organized structures typical of ecological or social systems (Figures 3, S3–S6). The seemingly weaker increase in *Max*[***S*_y_**] observed in the social-ecological networks is only an apparent effect of the scale. Indeed, as it will be shown later, suitable detection criteria of early warnings are more successful in mutualistic networks than in their random counterparts. Noise has the effect of amplifying the intensity of the warning sign (compare the scales in Figs. 2 and 3), while inducing weak random fluctuations with no substantial impact on the overall behavior of *Max*[***S*_y_**] at the onset of instability (see Supplementary Materials). In scale free networks the increase in *Max*[***S*_y_**] (Figure 3) is again only apparently muted. In fact, in these networks detection criteria are quite successful in recognizing early warning signs (Figure 4); moreover, local indicators (e.g., the variance of the most central node) can exhibit a more pronounced increase that can be used as an early warning sign of instability (Figure 1 and Supplementary Materials).

**Figure 3.**
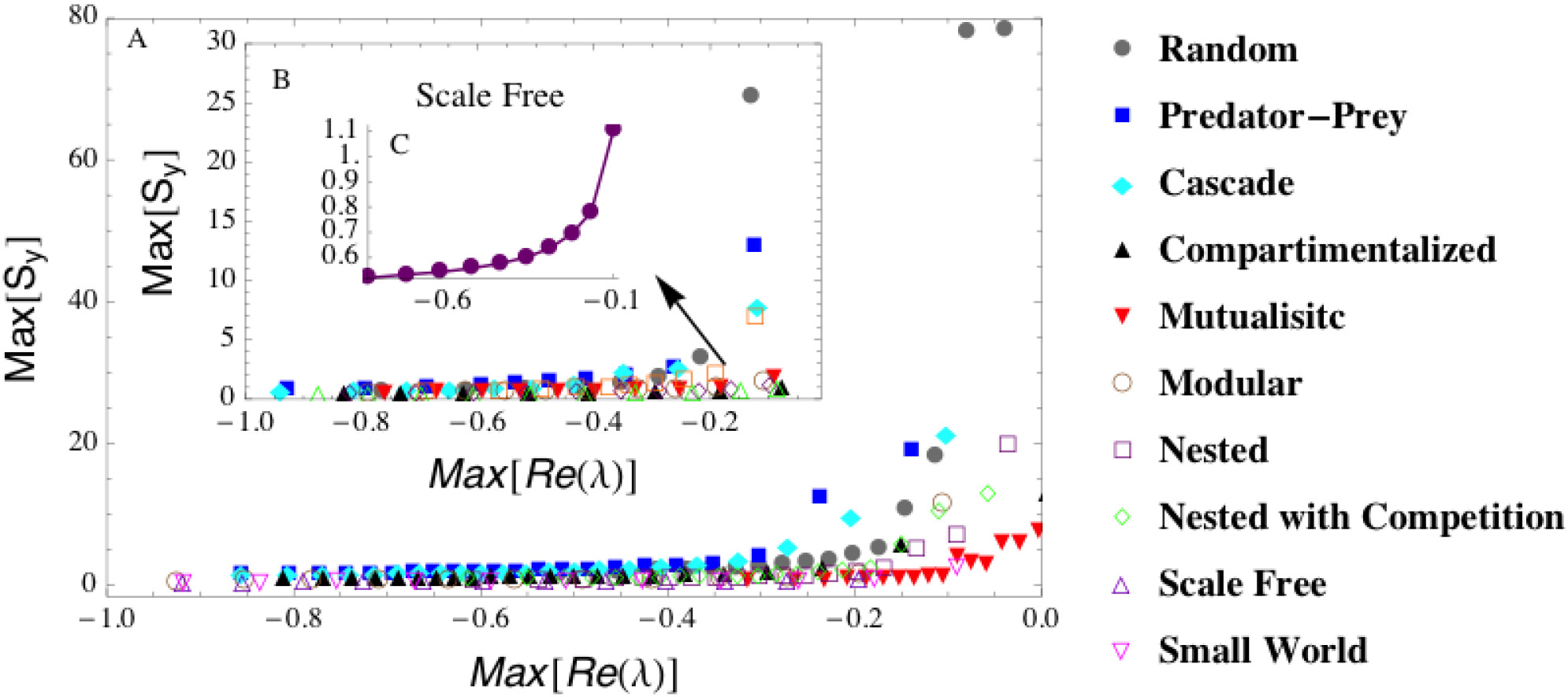
A) Case with random interaction strength (see methods). Main panel: instability is reached by increasing *p* (with *N* = 20; *C* = 0.2). First inset (B): *p* is constant while *C* increases between 0.1 and 1. C) Same as the first inset (B) but only for the scale-free network (notice the different scale on the vertical axis). The figures represent average behavior over 1000 realizations.

**Figure 4.**
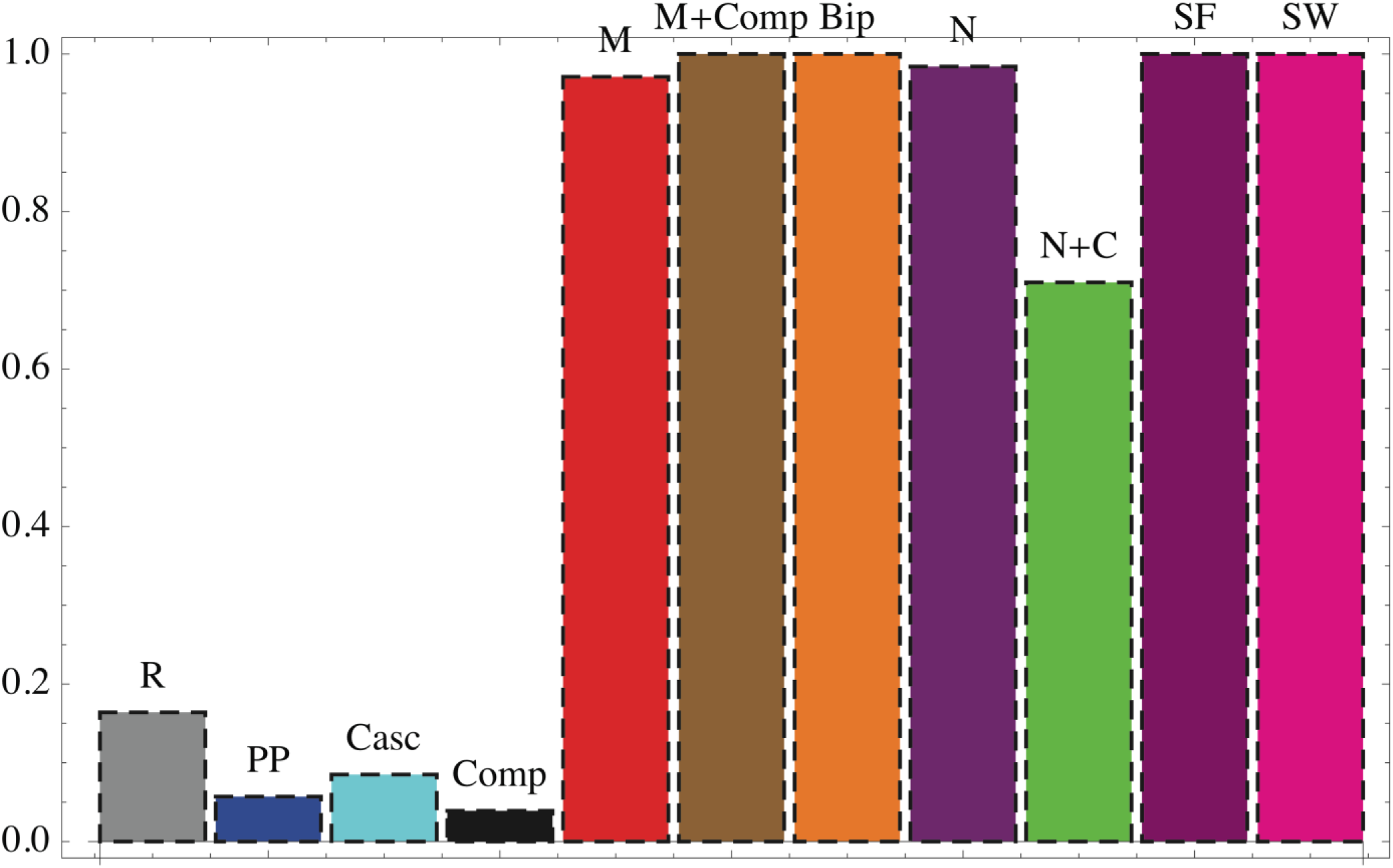
Distribution of the correlation, ρK, between *Max*[**S_y_**] and the parameter *p*, after 1000 realizations for the full disordered (not mean-field) case. If ρK is significant (p-value < 0.05) and ρ_K_ > 0.5 the increase in *Max*[**S_y_**] is interpreted as an early warning sign. We calculate these detection statistics for several realizations of each network structure and determine the probability of detecting the early warning sign of instability. We consider eleven different network architectures typical of ecological or social networks, including random (R), predator-prey (PP), cascade (Casc), compartmentalized (Comp), mutualistic (M), bipartite (Bip), nested (N), nested with competition (N + C), scale free (SF), and small world (SW). These networks have different structures for the adjacency matrix and different combination of interaction types, i.e (++) mutualistic, (+ −) antagonistic, (−−) competitive or a combination of them (See Supplementary Materials for more details).

### Conclusions

We have identified some suitable early warning signs in social-ecological networks in agreement with those identified by Ref. (33), and provided a theoretical framework for their interpretation. Overall, the performances of *Max*[***S*_y_**] as a leading indicator of instability change between random, antagonistic, mutualistic/social networks. This indicator gives an earlier and “sharper” warning sign in random than mutualistic and social networks. The warning sign, however, is harder to detect and is more likely to be missed in random and antagonistic networks than in their mutualistic or social counterparts (Figures 4, S16). Thus, by affecting the probability that early warnings are missed, the sign of the interactions within the network determines the consistency and reliability of this leading indicator. In fact, different realizations of the same network dynamics can yield different results in the behavior of *Max*[***S*_y_**] and thus this indicator might not detect in useful advance the emergence of instability (Figures S17–S18). The probability of true positives is close to 100% (i.e., negligible probability of false negatives) in mutualistic networks, and much smaller in random and antagonistic (predator-prey, cascade or compartment) networks (Figures 4, S16). Thus, while mutualistic networks are less stable than their antagonistic counterparts (17), their instability can be predicted with less uncertainty. An increase in *Max*[***S*_y_**], however, would not provide information on how close the system is to the onset of instability. Rather, it would just indicate that the system is losing resilience and approaching unstable conditions (29). Therefore, in contrast to previous expectations (16), it is not the heterogeneity in the topology of the network that plays a key role in the abruptness of critical transitions and our ability to predict them. Rather, it is the type of interactions between the nodes that determines how networks respond to external perturbations. In fact, there is a trade-off between local and systemic resilience: mutualism (++) is associated with a reduced local stability and resilience of the system (17,19), but does not induce abrupt critical transitions. In contrast, networks with mixtures of interaction types (+−,++, −−) exhibit shorter recovery times after displacement from equilibrium (i.e., a stronger local resilience) (17–18), but in these systems the emergence of systemic instability and critical transitions is more difficult to predict in useful advance.

This study combines stability theories from community ecology (2,17) to recent research on indicators of critical transition (7,9,16), and develops a unified framework that offers a new perspective for the evaluation of the resilience and anticipation of instability in social-ecological networks.

## Supplementary online materials

### S1. Structures of Socio-Ecological Interacting systems

a) *Random matrix* (1) with connectivity *C.* We pick an element A_i,j_ at random, and with probability *C* we assign a link between nodes *i-j* and *j-i* (A_j,i_). Each of these two links has a weight (or strength), *p*, that is positive (or negative) with probability 0.5; b) *Antagonistic matrix* (2), where connections are again random (with probability *C*), but if A_i,j_ has sign +(–), then A_j,i_ has sign – (+); c) *Cascade network* (3): links occur with a given probability, but form a hierarchical structure, whereby there is a top “predator” (with sign +) that feeds on all the other species (with sign −); then there is the second top predator, and so on; species left with the lowest ranking function as producers; d) *Compartmentalized* structure (5), formed by groups of antagonistic species that interact only with species within their own group; e) *Mutualistic network* (6), with a random matrix, but where both A_i,j_ and A_j,i_ have positive signs (++); f) *Modular mutualistic* matrix (5): nodes are divided into communities that positively interact within their groups (++). g) *Bipartite Mutualistic* matrix (6): nodes are divided into two groups, and each node positively interacts only with nodes of the other group (++); h) *Bipartite Nested Mutualistic* (7): a bipartite graph is generated with hierarchical structure where specialist nodes (i.e., with only few mutualistic links) tend to interact with a suitable subset of the mutualistic partners of the generalist nodes (++); i) *Nested Mutualistic with Competition* (8): nodes are divided into two groups; each node positively interact with nodes of the opposite group (++), while compete with nodes of the same group (−−); j) *Barabasi-Albert (BA) networks* (9): binary one-zero networks are generated with power-law degree distribution. This algorithm simulates a preferential attachment process, in which a new vertex with *d* edges is added at each step. The BA graph displays a scale-free behavior that strongly correlates with the network's robustness to failure. We then assign to each link a positive weight *p*; The connectivity of the BA networks is controlled indirectly by the parameter *d*. In our simulation we set *d* = 1 to generate low connectivity networks, *d* = 2 for average connectivity and *d* = 4 for high connectivity. k) *Watts-Strogatz (WS) network* (10): this is a one-parameter (degree of disorder r) model that interpolates between an ordered finite dimensional lattice and a random graph. Its main property is to display both high clustering coefficient and small world property. The WS model displays this duality for a wide range of the rewiring probabilities r. In our simulation we used r = 0.3. The connectivity of the WS model depends on the starting 2k-regular graph (10). For low connectivity we set k = 2; for average connectivity k = 4; for high connectivity k > 5.

The linkage density in each network is described by the parameter *C* (the connectivity).

We generate each of the structures (*a*)-(*k*) keeping constant the connectivity *C* and changing the strength of the interactions (alternatively, we fix *p* and vary *C*), until the dynamics become unstable (1,6). Thus, we use the parameter, *p*, as an indicator of the weights of the interactions between connected nodes (connectivity). We consider three different levels of disorder in the control parameter *p*: 1) *Mean Field:* where we assign a constant value, *p*. We do not consider antagonistic mean field networks (i.e., with constant |*p*| but (+−) interactions) because they are always stable (see below); 2) *Weak disorder*: the interaction strength is drawn from a normal distribution with mean *p* and standard deviation 0.1 *p* 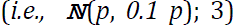 *Strong disorder*: interaction strengths are drawn at random from 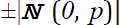 (6). Therefore, in all these cases, by increasing the value of *p* (*C*) the resilience decreases until the system become unstable for *p* = *p*_*c*_, or *C* = *C*_*c*_ (6).

### S2.1 Early Warnings in Multidimensional Mean Field Networks

In the mean field case with (++) interactions, the matrix is symmetric (**A** = **A**^T^) and therefore all eigenvalues are real numbers. In this case we note that, if **U** is the matrix of the eigenvectors of **A** (i.e. **A u**_i_= λ_i_ **u**_i_), then **U A U**^T^=**diag**(**λ**), where **diag**(**λ**) is a diagonal matrix with all the eigenvalues of **A**. Therefore in this case we can write Eq. (2) in the main text as:

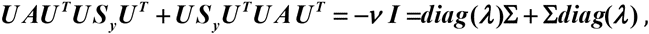

where ***UU***^T^ **= *I*** and **Σ = *U S U***^T^ is a diagonal matrix whose *i*-th eigenvalue is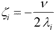

Therefore if the socio-ecological system reaches an instability (i.e. λ_i_=0) for a given *i*, then there is at least one element of the matrix **Σ** and thus of **S_y_** that diverges.

We also notice that in the mean field case we omit the analysis of early-warning performed on antagonistic (+−) matrices, **A′**. In fact, if **A′** is anti-symmetric (i.e., **A′**=-**A′**^T^, and thus with predator prey, cascade or compartmentalized interactions), then the early warning analysis on those matrices is trivial as their stability does not depend on *p* or *C*. In fact, if we multiply the off-diagonal terms of those matrices by the imaginary unit *i*, i.e. **B** = *i* (**A**′-**I A′)**, then **B** is Hermitian and has all real eigenvalues. Therefore **A**′-**I A′** has all pure complex eigenvalues independently of *p* and *C*, which means that **A′** is always stable – given that the self interaction terms are negative constant, i.e. A′_i,i_=−d < 0.

Figures S1 and S2 show a rise in *Max*[**S_y_**] as the system approaches instability in mean field networks.

**Figure S1.**
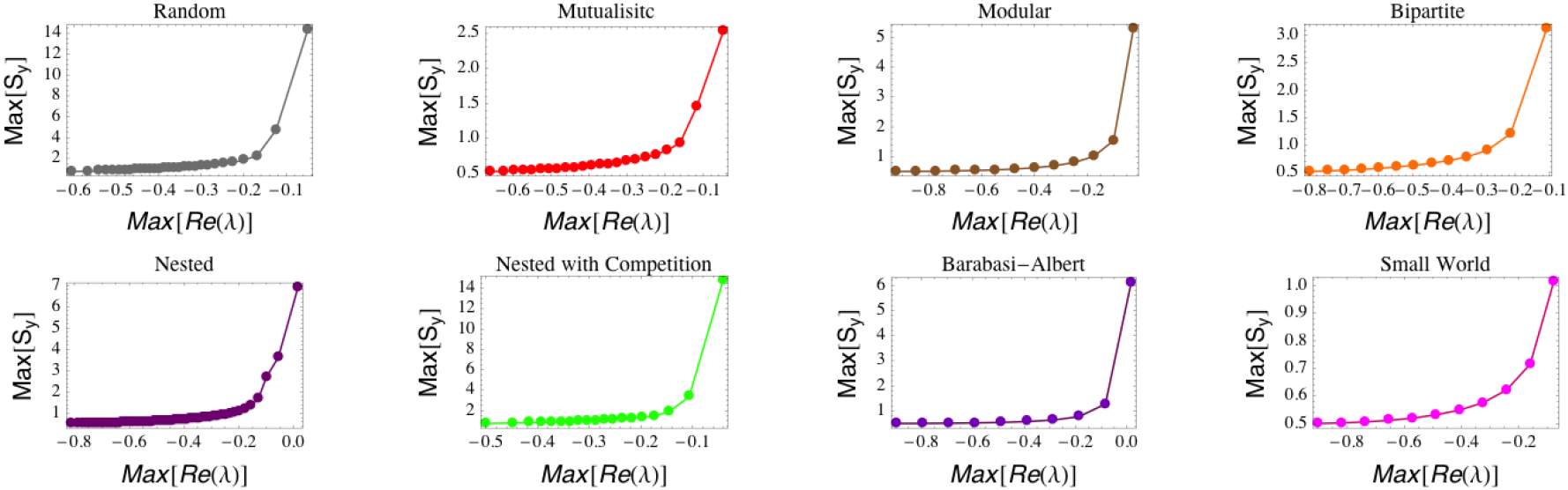
Increase in *Max*[**S_y_**] as *Max*[Re(λ)] tends to zero for mean field networks of size, *N* = 20, C = 0.2. Increasing values of *Max*[Re(λ)] are obtained by increasing the interaction strength, *p.* The plotted values are the ensemble averages of 1000 realizations.

**Figure S2.**
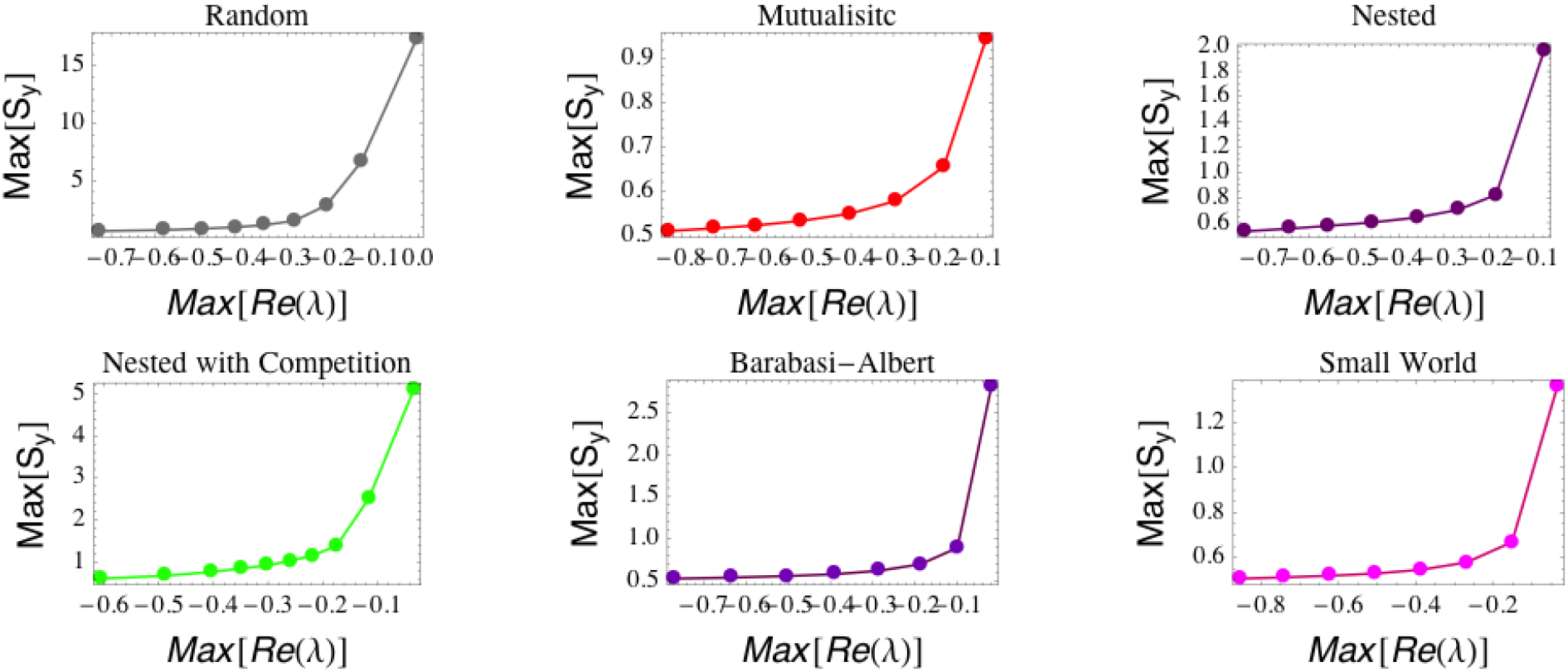
Increase in *Max*[**S_y_**] as *Max*[Re(λ)]→0 for mean field networks of size, *N* = 20, p<<p_c_. Increasing values of *Max*[Re(λ)] are obtained by increasing the connectivity, *C.* The plotted values are the ensemble averages of 1000 realizations.

**Figure S3.**
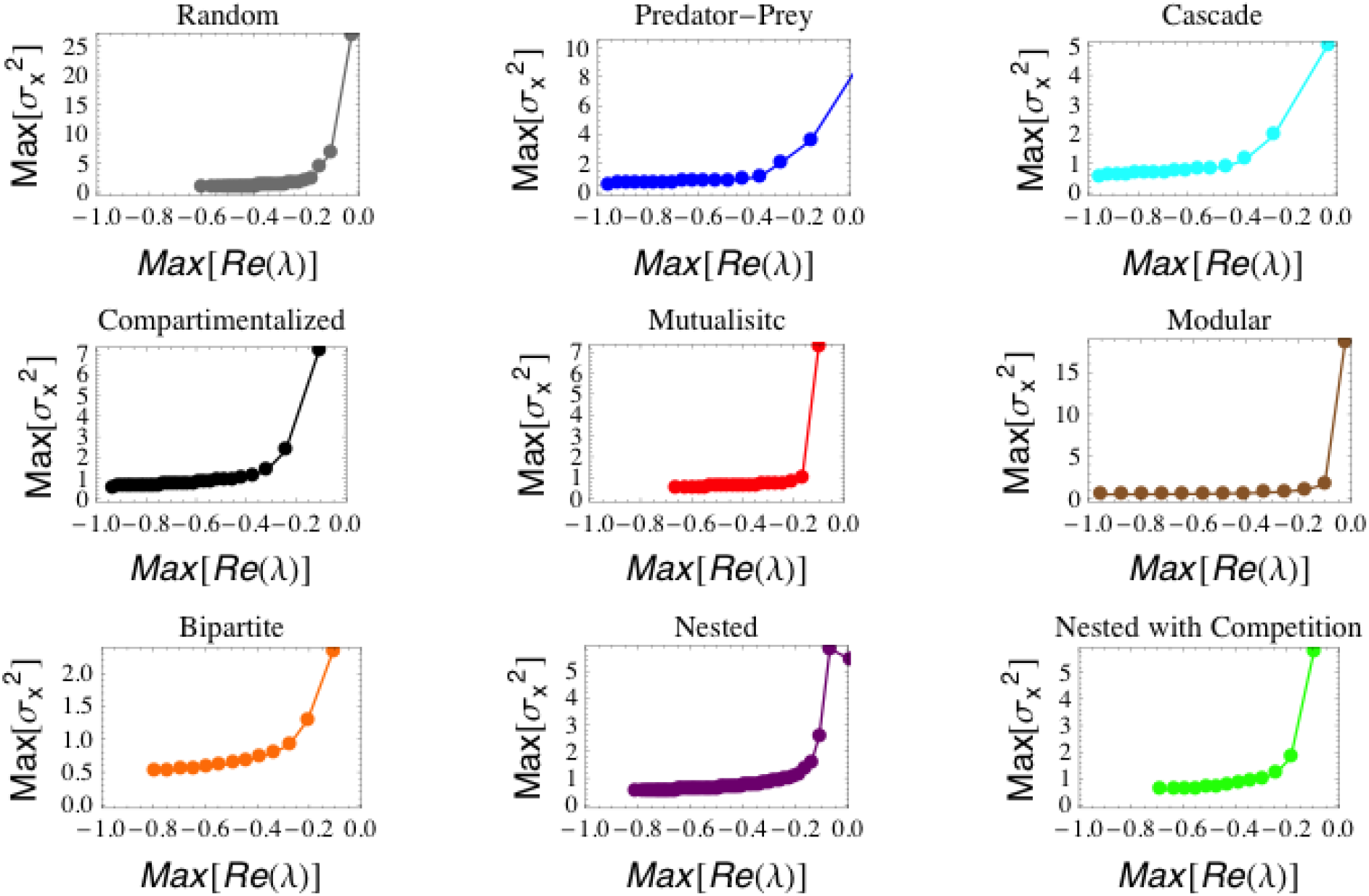
Increase in *Max*[**S_y_**] as *Max*[Re(λ)] tends to zoro for complex networks with “weak” disorder (see Section 1) of size, *N* = 20 and C = 0.2. Increasing values of *Max*[Re(λ)] are obtained by increasing the interaction strength, *p.* The plotted values are the ensemble averages of 1000 realizations.

**Figure S4.**
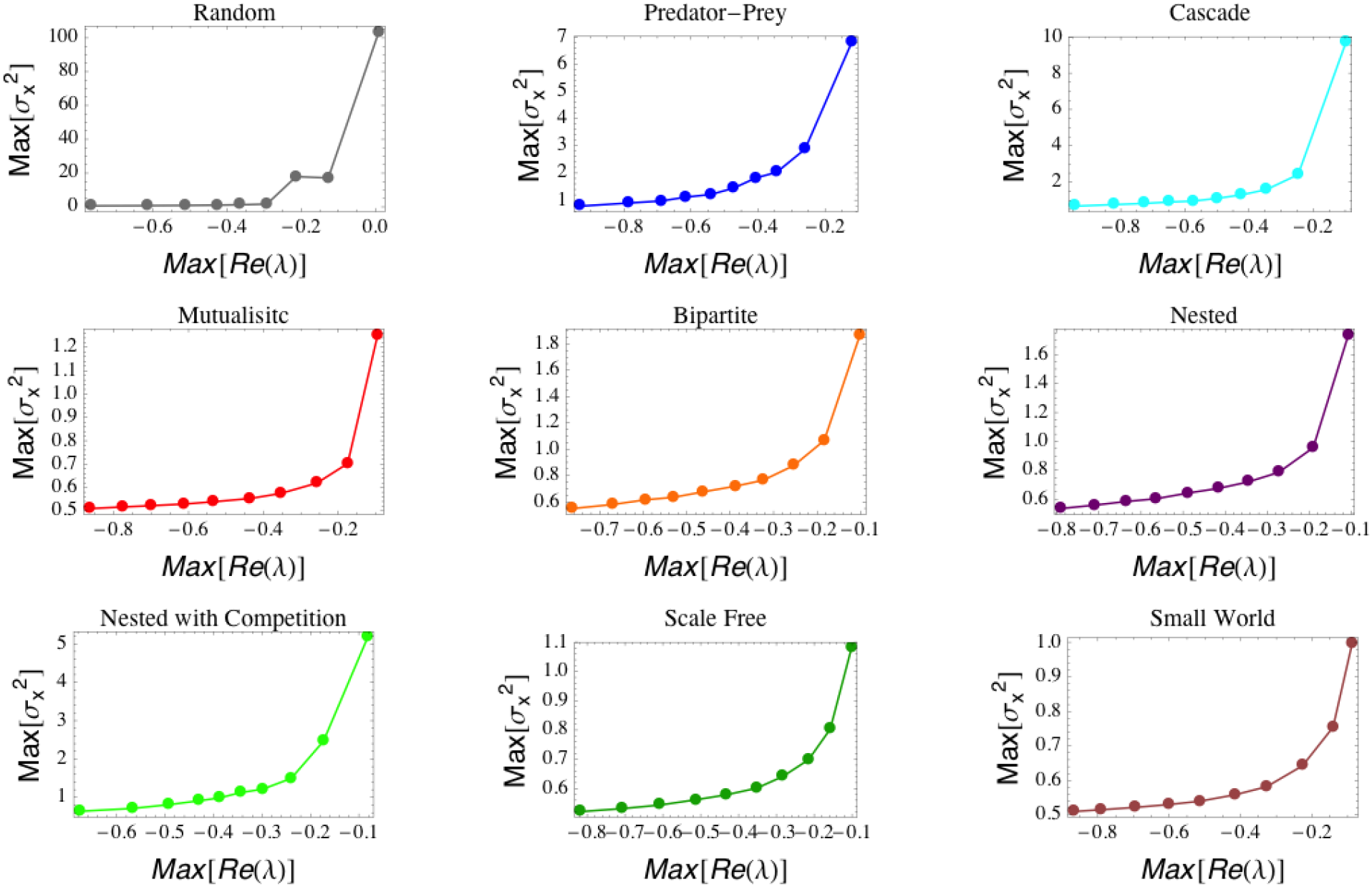
Increase in *Max*[**S_y_**] as *Max*[Re(λ)]→0 for complex networks with “weak” disorder (see Section 1) of size, *N* = 20 and p<<p*_c_*. Increasing values of *Max*[Re(λ)] are obtained by increasing the connectivity, *C.* The plotted values are the ensemble averages of 1000 realizations.

**Figure S5.**
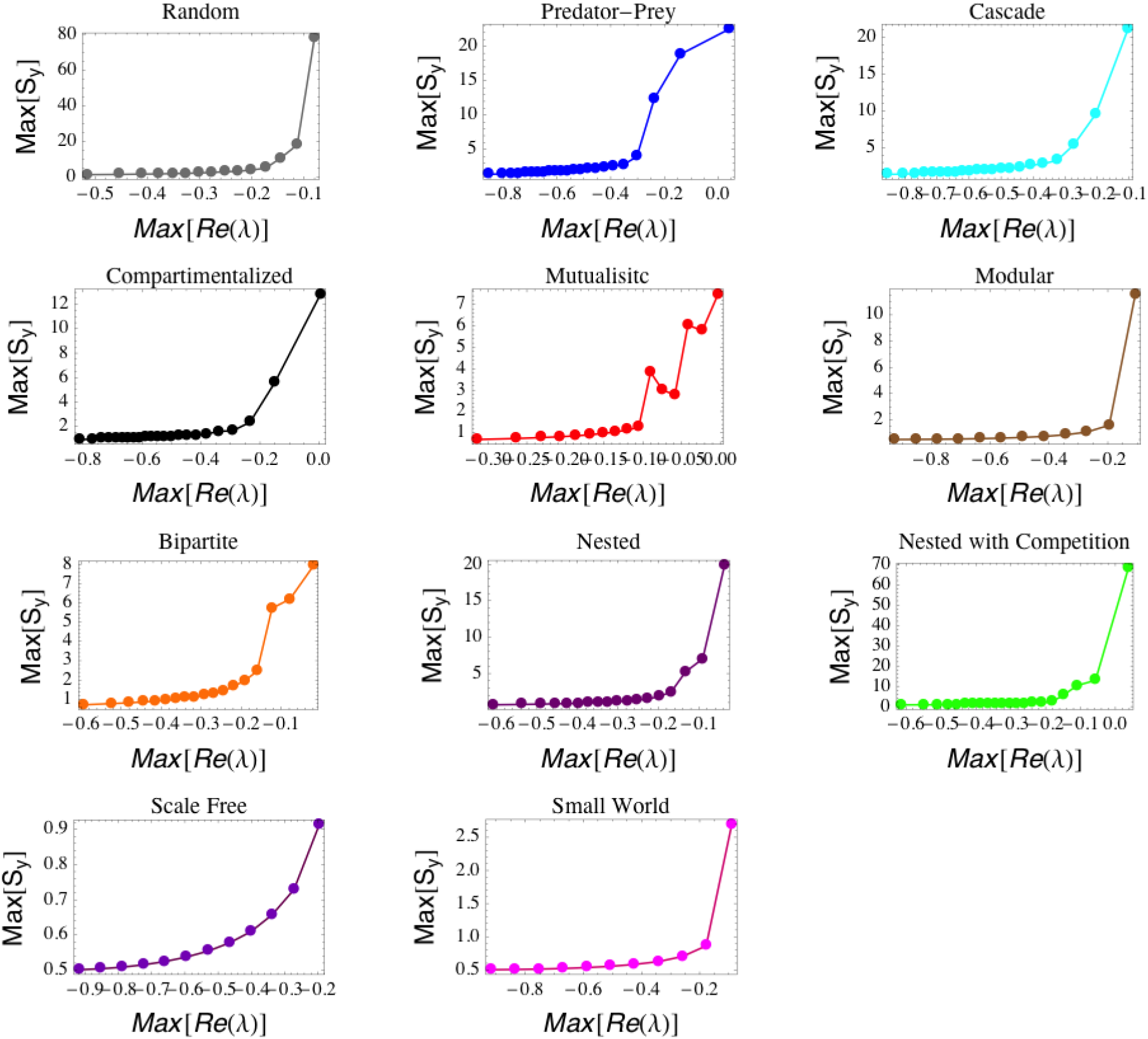
Increase in *Max*[**S_y_**] as *Max*[Re(λ)] tends to zero for complex networks with “strong” disorder (see Section 1) of size, *N* = 20 and C = 0.2. Increasing values of *Max*[Re(λ)] are obtained by increasing the interaction strength, *p.* The plotted values are the ensemble averages of 1000 realizations.

**Figure S6.**
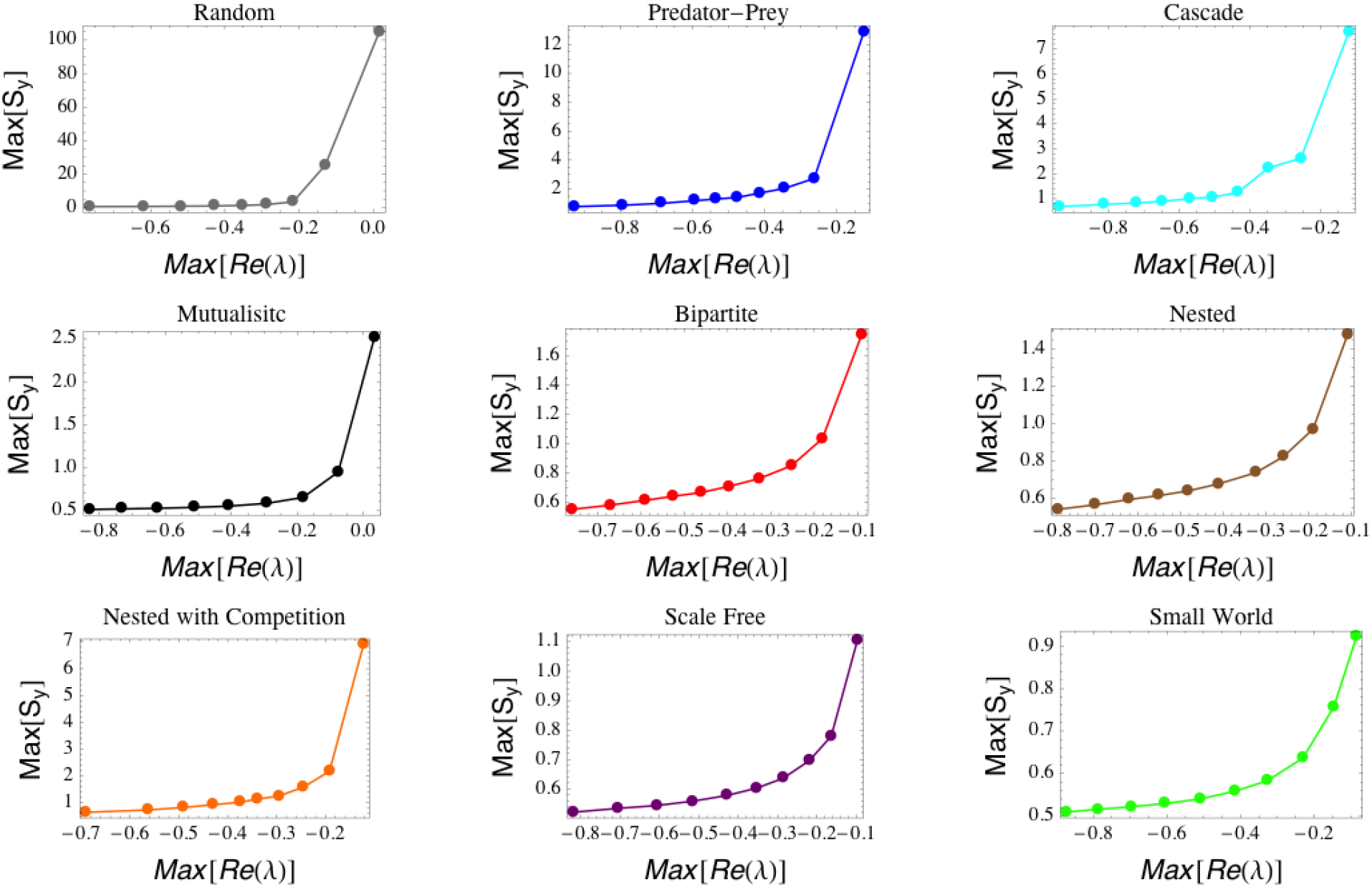
Increase in *Max*[**S_y_**] as *Max*[Re(λ)] tends to zero for complex networks with “strong” disorder (see Section 1) of size, *N* = 20. Increasing values of *Max*[Re(λ)] are obtained by increasing the connectivity, *C.* The plotted values are the ensemble averages of 1000 realizations.

### S3. Indicators and Node Properties

In this section we show more analysis on the effectiveness of different node variances as precursors of instability.

**Figure S7.**
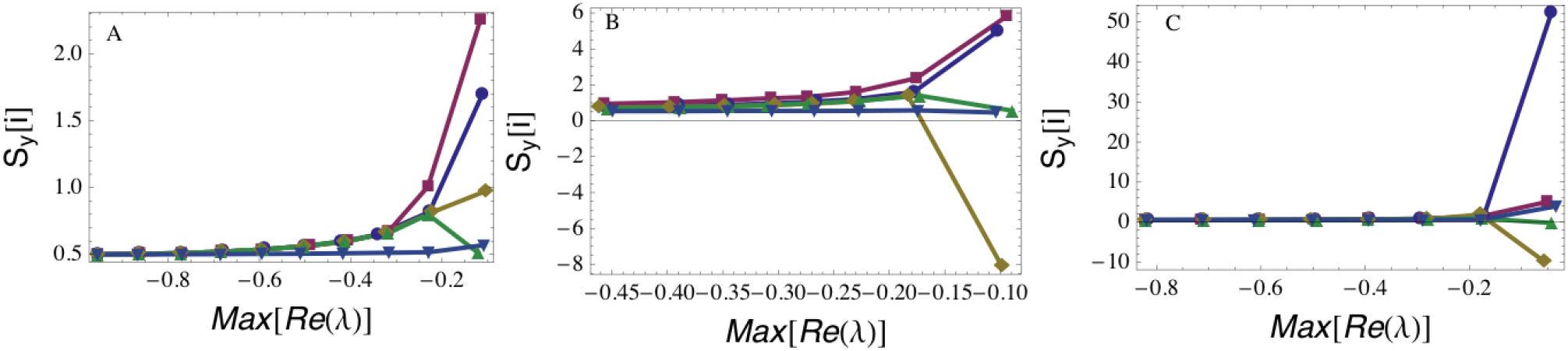
Elements of the covariance matrix **S_y_** corresponding to nodes with the highest number of connections (green), lowest number of connections (light blue), highest eigenvector centrality (gold), max[**S_y_**] (violet) and max[**S_y_**]-min[**S_y_**] (purple), in the case of: (A) mutualistic, (B) mutualistic nested with competition, (C) small world interactions, for mean field networks (of size *N* = 20 and connectivity *C* = 0.3). The plotted values are the ensemble averages of 1000 realizations.

**Figure S8.**
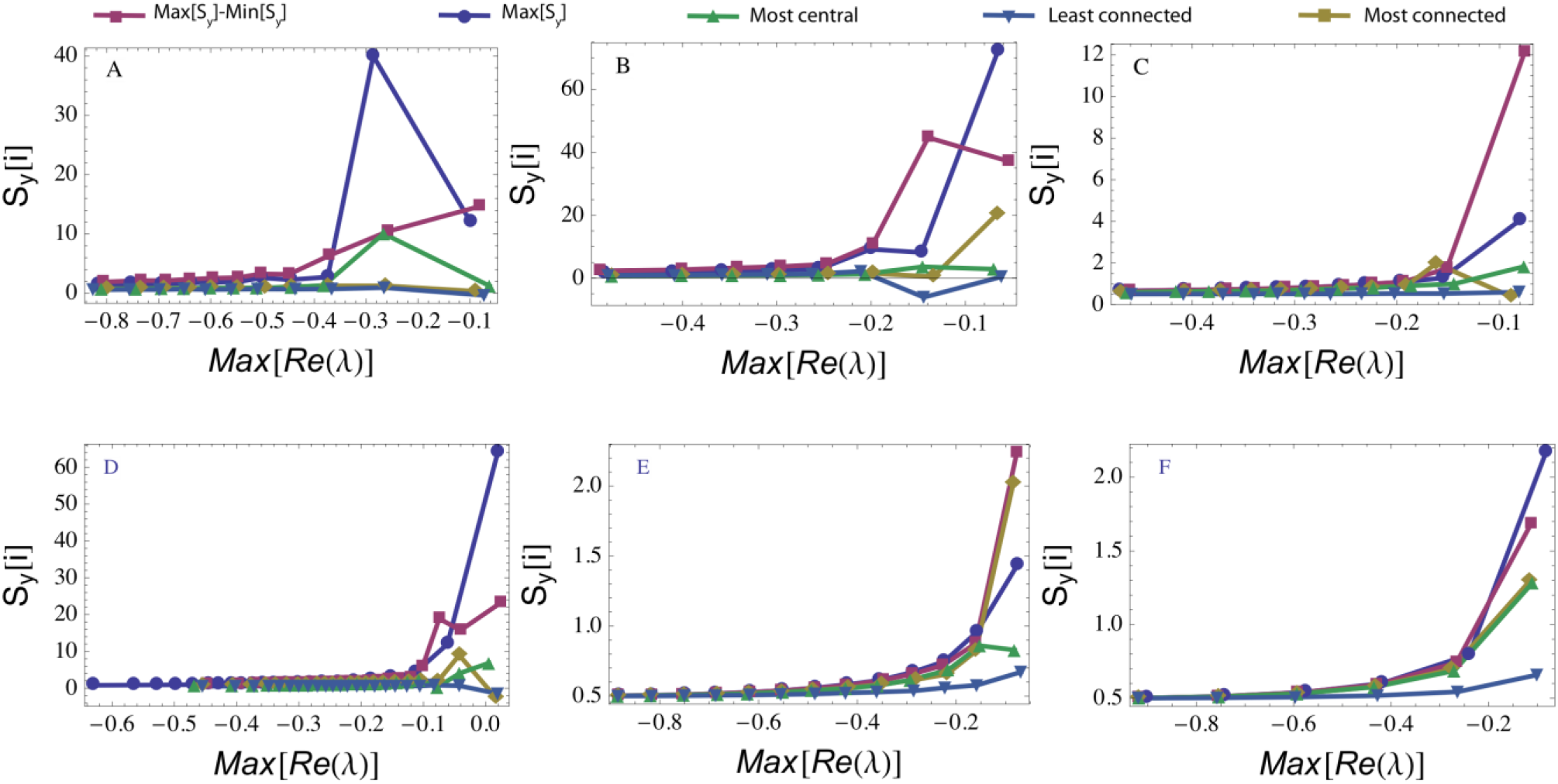
Elements of the covariance matrix **S_y_** corresponding to nodes with the highest number of connections (green), lowest number of connections (light blue), highest eigenvector centrality (gold), max[**S_y_**] (violet) and max[**S_y_**]-min[**S_y_**] (purple) in the case of (A) random, (B) predator-prey, (C) mutualistic, (D) mutualistic nested with competition, (E) Small world, (F) Barabasi-Albert, networks with strong disorder (of size *N* = 20 and connectivity *C* = 0.3). The plotted values are the ensemble averages of 1000 realizations.

**Figure S9.**
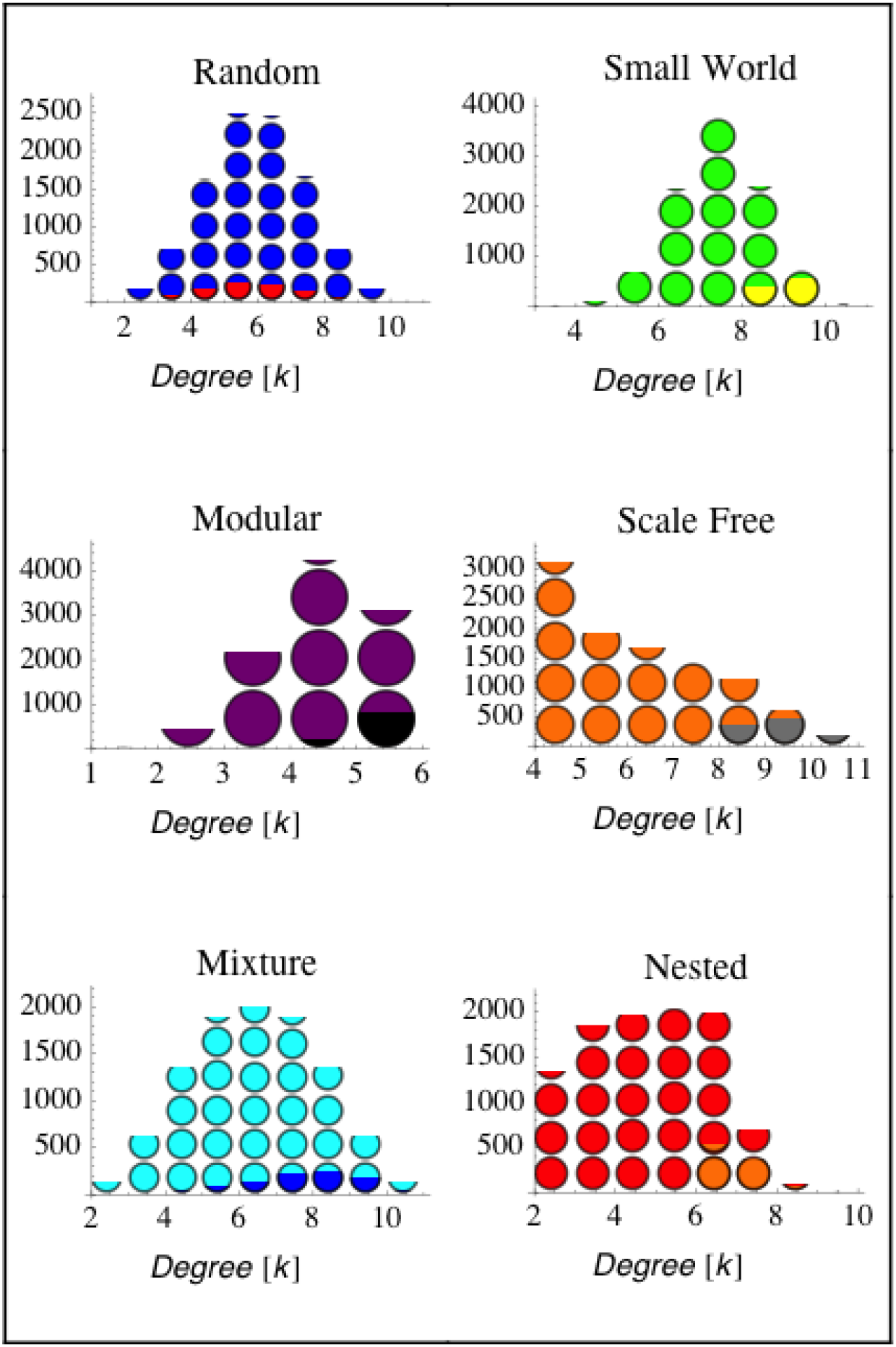
Frequency distribution of the degrees (i.e., number of connections) of the networks’ nodes and (with partially filled circles) of the nodes associated with the maximum value of the covariance matrix **S_y_** in mean field networks with a variety of interactions. Based on a set of 100 realizations. Notice how, in mutualistic networks the node corresponding to max[**S_y_**] is associated with the nodes with the highest degrees (i.e. the generalist species).

**Figure S10.**
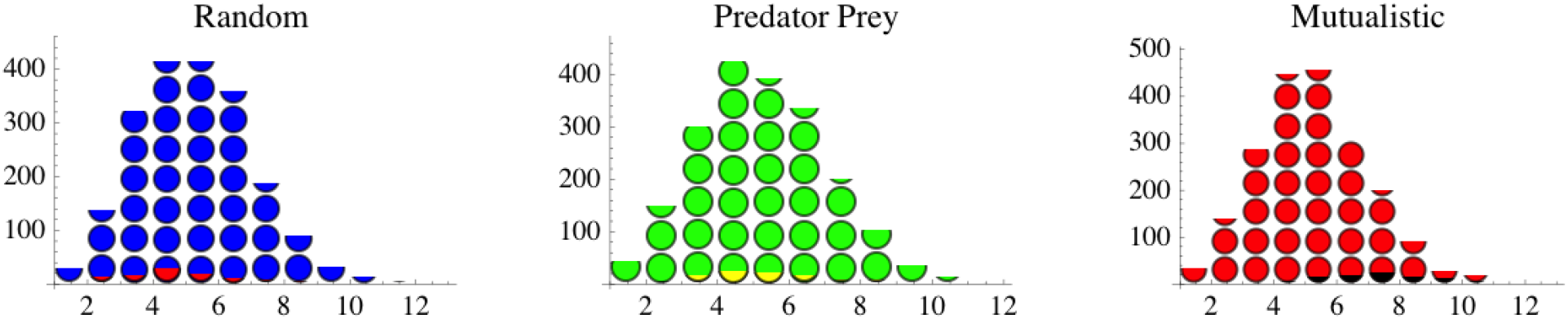
Frequency distribution of the degrees (i.e., number of connections) of the networks’ nodes, and (with partially filled circles) frequency distribution of the nodes associated with the maximum value of the covariance matrix **S_y_** in “strongly” disorganized networks with a variety of interactions. Based on a set of 100 realizations. Notice how, in mutualistic networks the node corresponding to max[**S_y_**] is associated with the nodes with the highest degrees (i.e. the generalist species).

**Figure S11.**
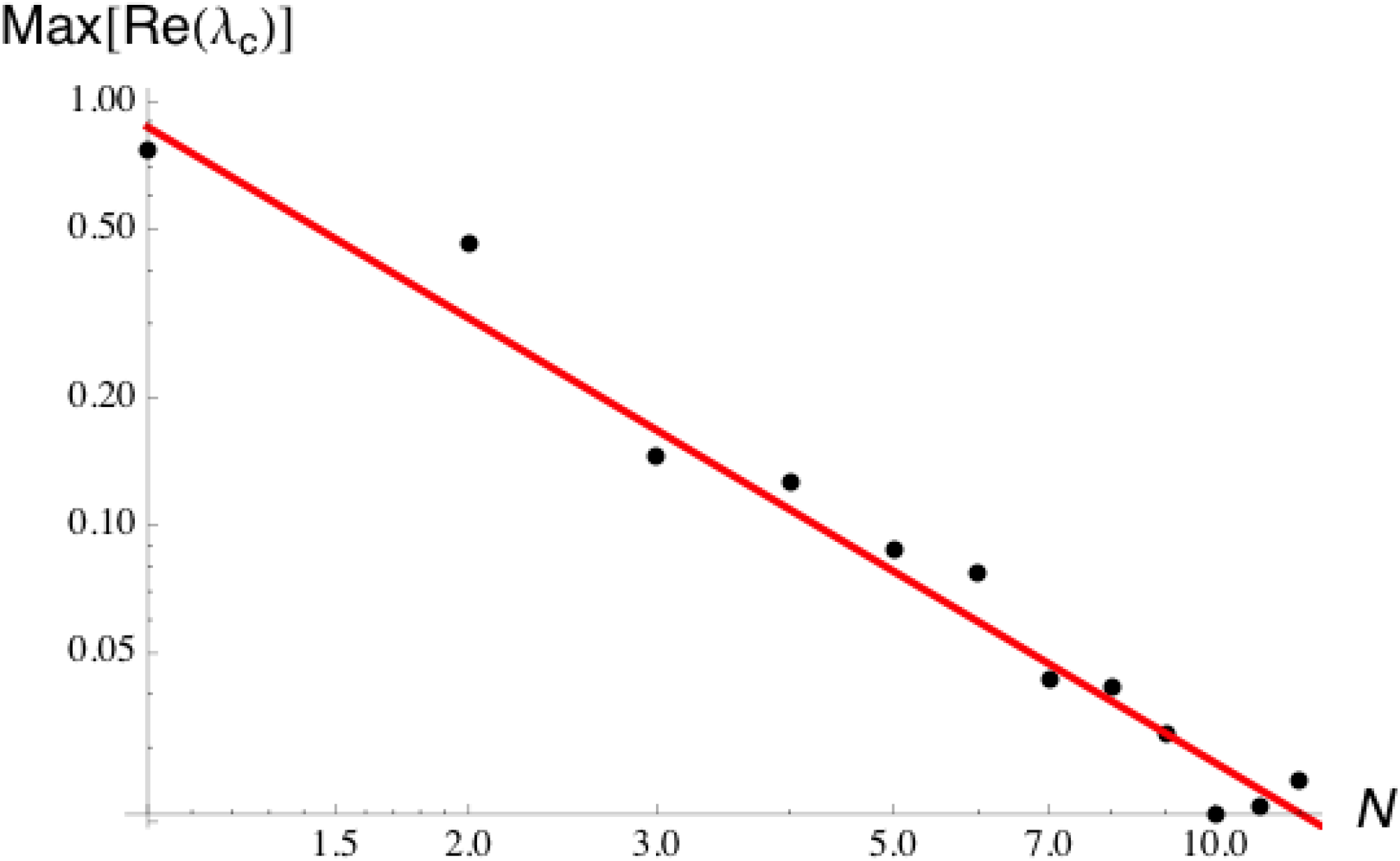
Effect of the network size on the magnitude of the early warning sign. Maximum real part (in absolute value) of the network's eigenvalues as a function of the network size, *N* for a random network with C = 0.25 and *p* = *p_c_*=1/√NC. As *N* increases the max of Re(λ) tends to zero as *Max*[Re(λ)] ∼ N^−1.5^and the resilience of the system decreases, while the “height” of the early warning increases. Therefore, as *N* increases, the early warning sign becomes sharper (see also Figure 1 in the main text).

### S4. Early Warning for Time Correlation Matrix and Power Spectrum

We calculate the time lag correlation matrix, **ρ**_y_(Δ) (where Δ is the time lag), and the power spectrum, ***P***_y_(ω) (where ω is the frequency) for the steady state dynamics:

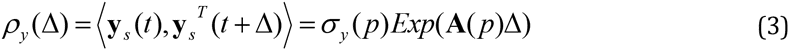

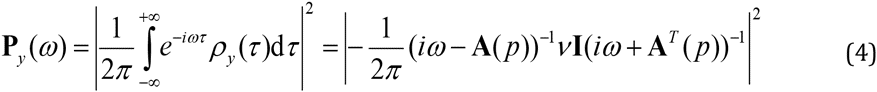

The following figures show the behavior of the autocorrelation and power spectrum as the system approaches instability.

**Figure S12.**
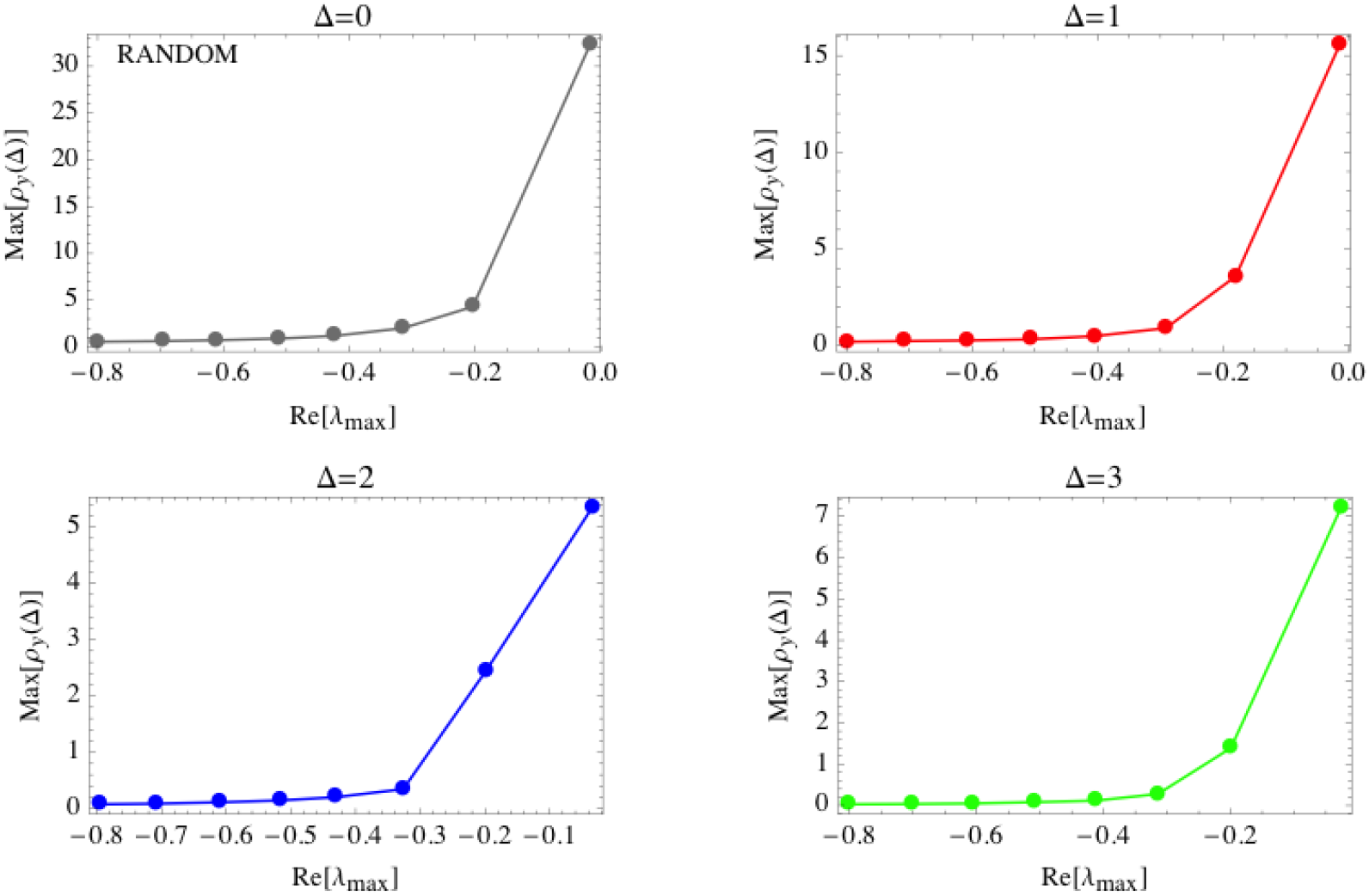
Increase in *Max*[**ρ_y_**] as *Max*[Re(λ)] tends to zero for “strongly” disordered networks with a random architecture with *N* = 20 and *C* = 0.3. Increasing values of *Max*[Re(λ)] are obtained by increasing the interaction strength, *p.* The plotted values are ensemble averages of 100 realizations.

**Figure S13.**
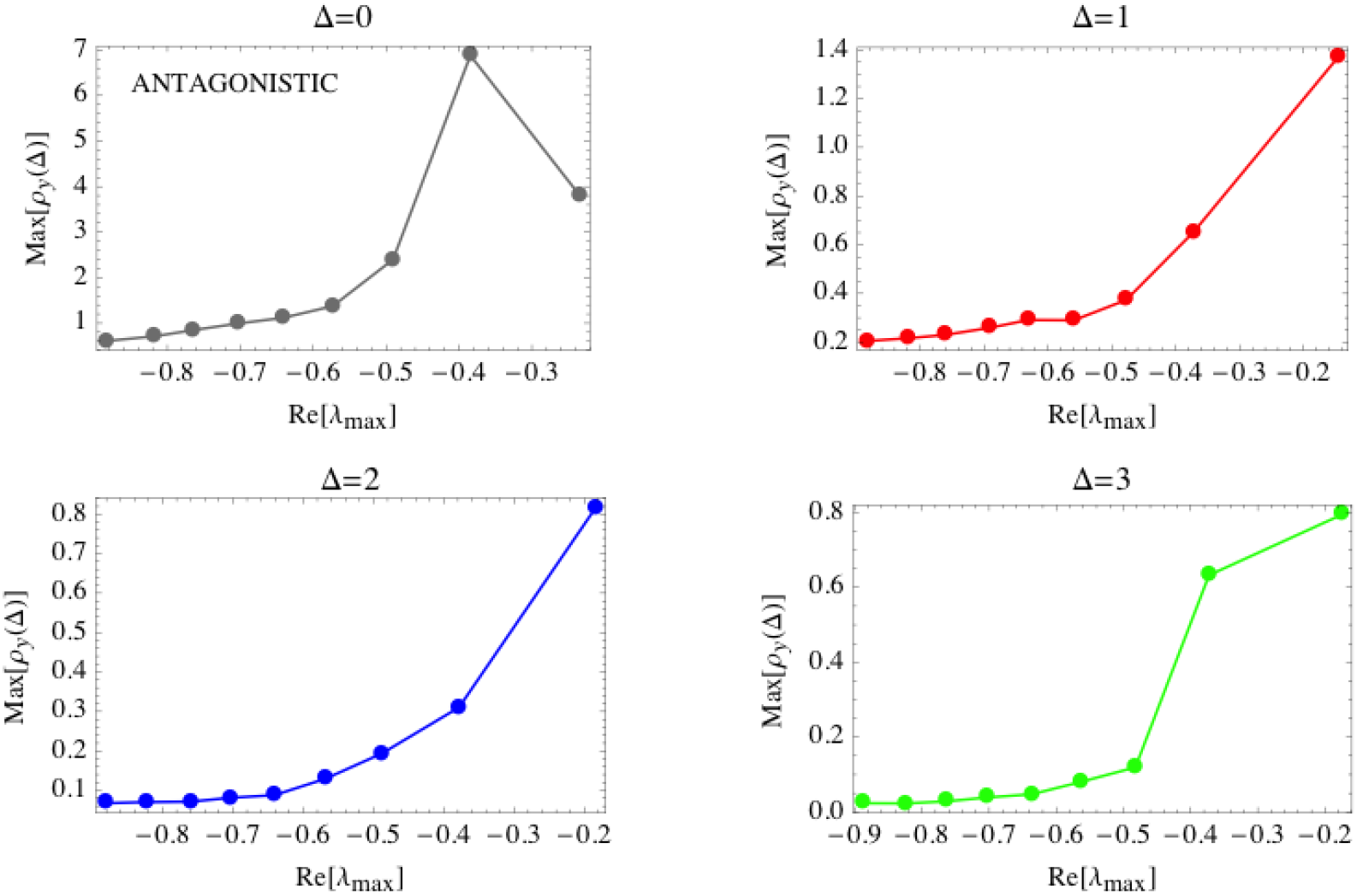
Increase in *Max*[**ρ_y_**] as *Max*[Re(λ)] tends to zero for “strongly” disordered networks with a predator-prey architecture and *N* = 20, *C* = 0.3. Increasing values of *Max*[Re(λ)] are obtained by increasing the interaction strength, *p.* The plotted values are ensemble averages of 100 realizations.

**Figure S14.**
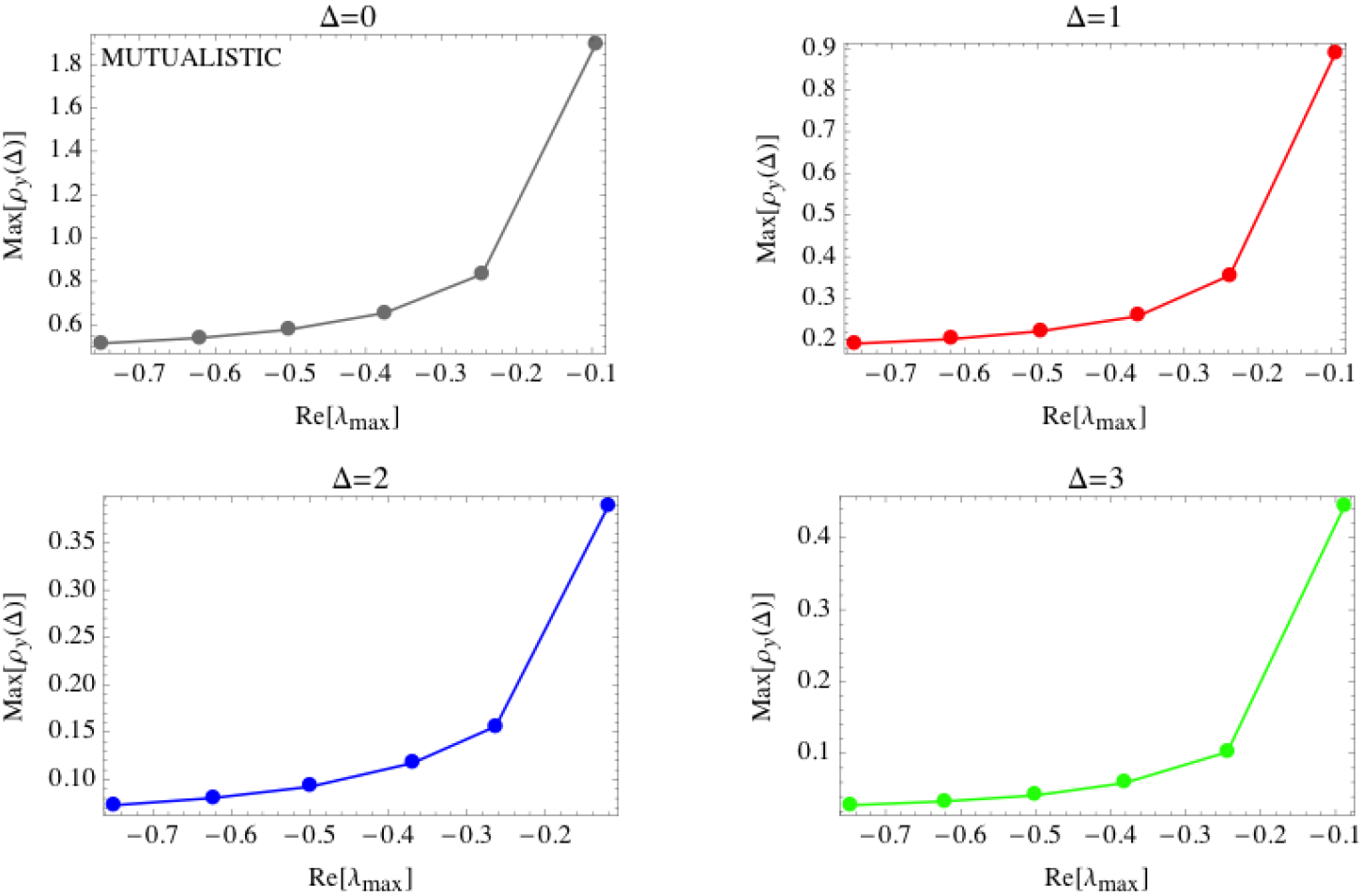
Increase in *Max*[**ρ_y_**] as *Max*[Re(λ)] tends to zero for “strongly” disordered networks with a mutualistic architecture and *N* = 20, *C* = 0.3. Increasing values of *Max*[Re(λ)] are obtained by increasing the interaction strength, *p.* The plotted values are ensemble averages of 100 realizations.

**Figure S15.**
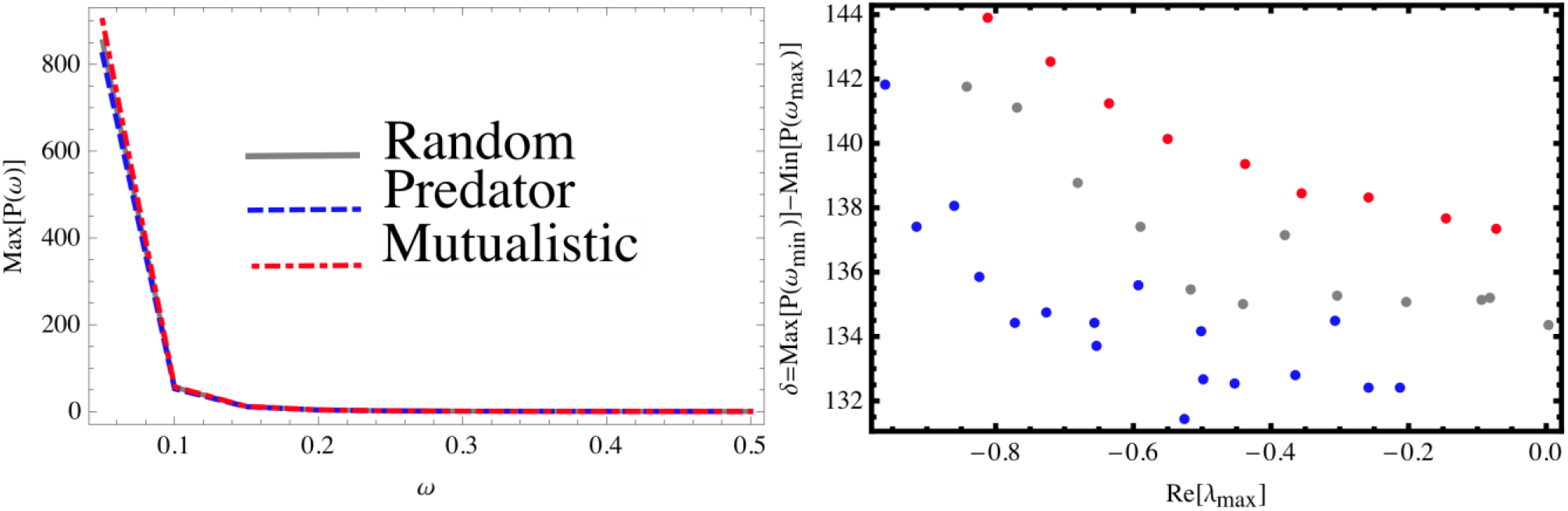
Left panel: max element of the power spectrum matrix as a function of frequency for three different architectures. The impact of the structure on the spectrum is negligible. Right panel: Power spectrum evaluated in the minimum and maximum frequency as *p* tends to *p_c_* (and thus *Max*[Re(λ)] tends to zero) for strongly disordered systems (*N* = 20 and C = 0.2) with random, predator-prey, and mutualistic interactions. Increasing values of *Max*[Re(λ)] lead to a decrease in δ= *Max*[P(ω_min_)] - *Max*[P(ω_max_)], that therefore might be considered a precursor for a critical transition. However, the intensity of this early warning sign is quite weak and thus difficult to detect. The plotted values are ensemble averages of 100 realizations.

### S5. Early Warning Detection

To evaluate whether the onset to instability can be anticipated in time by an increase in *Max*[**S_y_**] (or in other suitably chosen elements of **S_y_**), we test the correlation (11) between *Max*[**S_y_**] and the control parameter (*p* or *C*) that is gradually varied to increase *Max*[Re(λ)] up to a given threshold (here chosen equal to −0.2). If the correlation, ρ_k_, (evaluated with the Kendall-τ test) is significant and greater than 0.5, the increase in *Max*[**S_y_**] is interpreted as an early warning sign. We repeat this analysis for 1000 realizations of the random interaction strength network and determine the distribution of correlations along with the number of realizations with positive warning sign.

**Figure S16.**
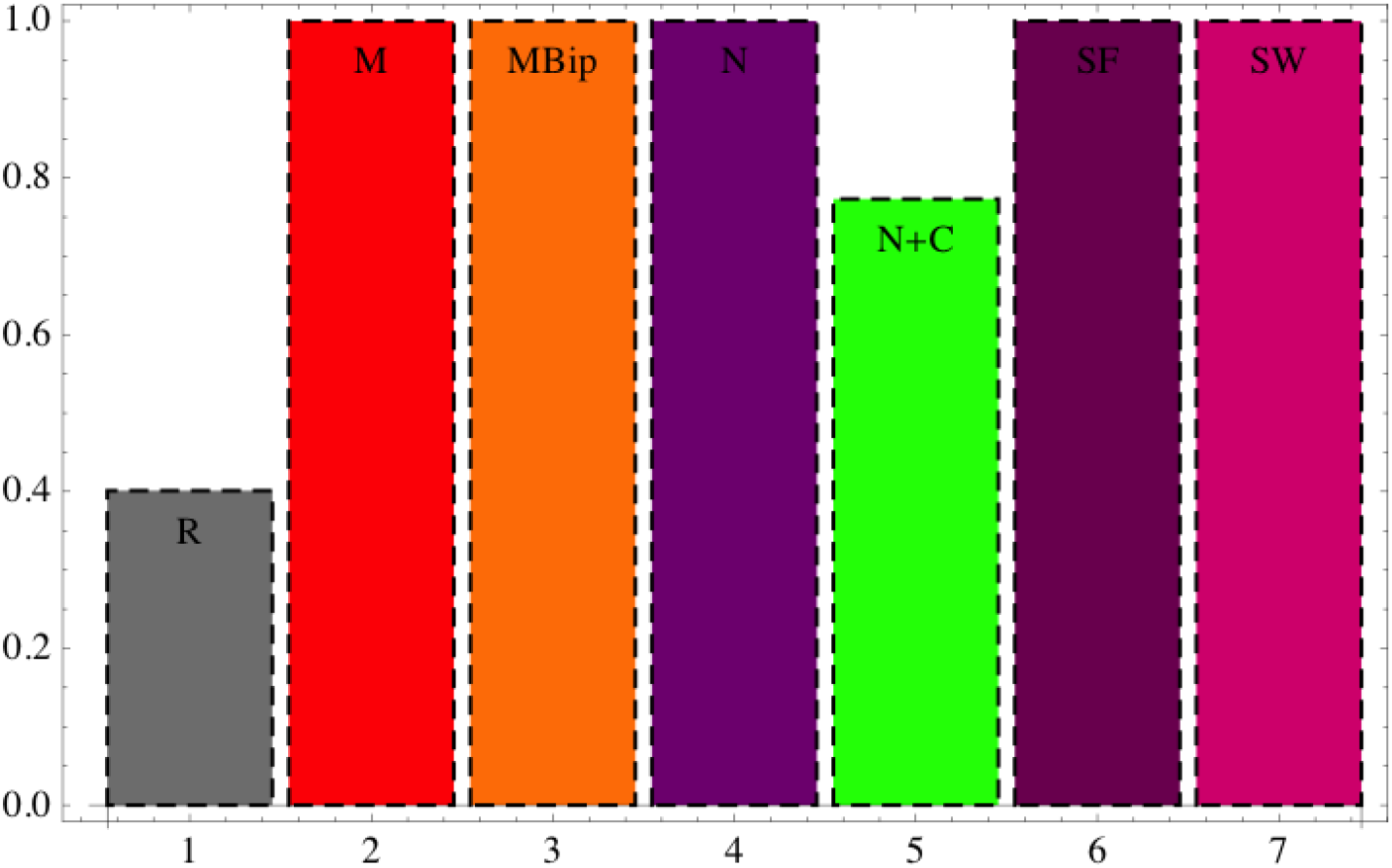
Probability of detecting true positives (i.e. of not missing a warning sign) in the case of mean field networks, using the same detection criteria as in Figure 4.

**Figure S17.**
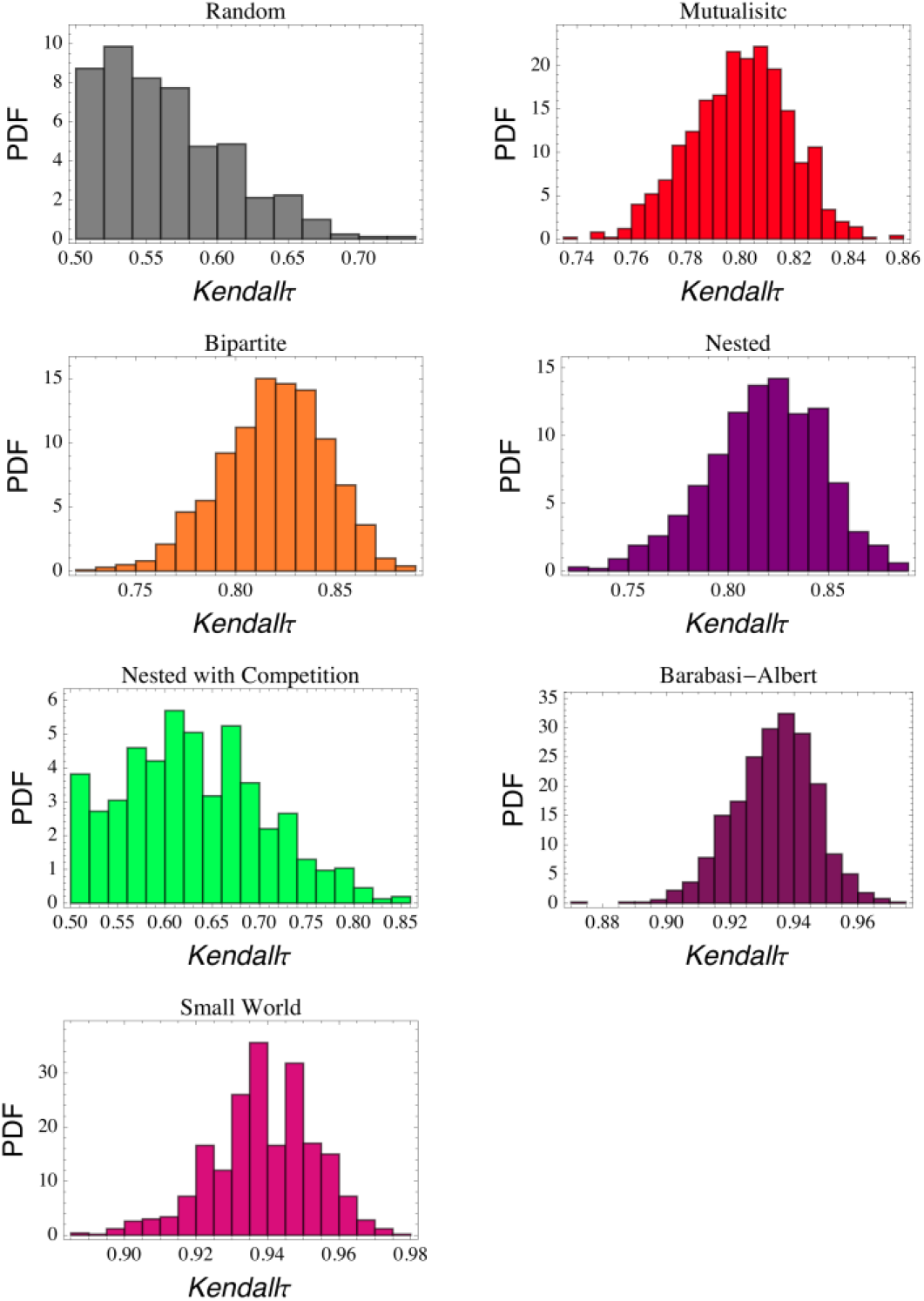
Frequency distribution of the ρ_K_ statictics used to detect early warning signs of instability in the case of mean field networks.

**Figure S18.**
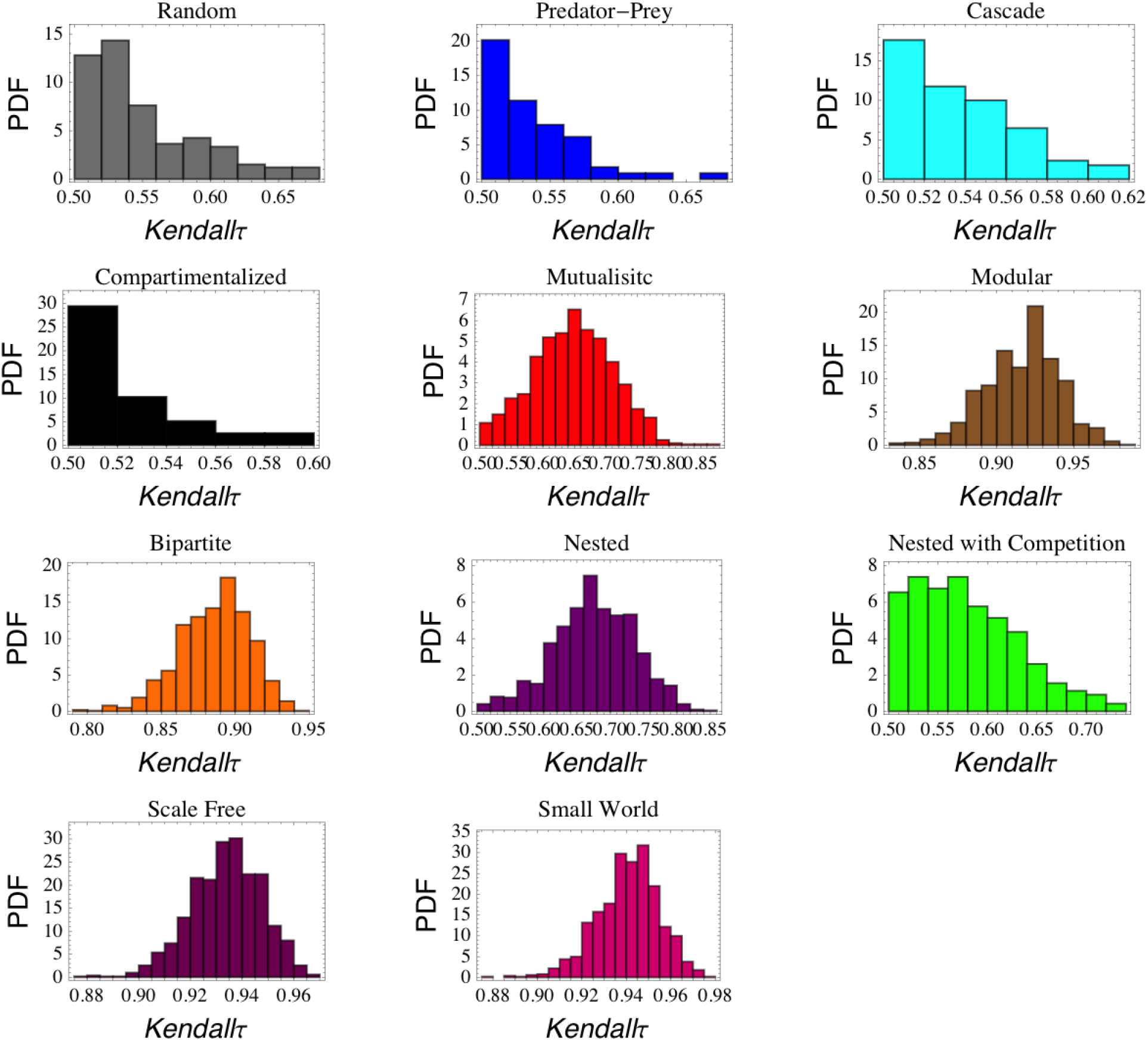
Frequency distribution of the ρ_K_ statictics used to detect early warning signs of instability in the case of full disordered networks.

